# GENERATION OF FUNCTIONAL NEURONS FROM ADULT HUMAN MUCOSAL OLFACTORY ENSHEATHING GLIA BY DIRECT LINEAGE CONVERSION

**DOI:** 10.1101/2023.09.28.559940

**Authors:** María Portela-Lomba, Diana Simón, Marta Callejo-Móstoles, Gemma de la Fuente, David Fernández de Sevilla, Vega García-Escudero, M. Teresa Moreno-Flores, Javier Sierra

## Abstract

A recent approach to promote central nervous system (CNS) regeneration after injury or disease is direct conversion of somatic cells to neurons. This is achieved by transduction of viral vectors that express neurogenic transcription factors. In this work we propose adult human mucosal olfactory ensheathing glia (hmOEG) as a candidate for direct reprogramming to neurons due to its accessibility and to its well-characterized neuro-regenerative capacity. After induction of hmOEG with the single neurogenic transcription factor NEUROD1, the cells under study exhibited morphological and immunolabeling neuronal features, fired action potentials and expressed glutamatergic and GABAergic markers. In addition, after engraftment of transduced hmOEG cells in mouse hippocampus, these cells showed specific neuronal labeling. Thereby, if we add to the neuro-regenerative capacity of hmOEG cultures the conversion to neurons of a fraction of their population through reprogramming techniques, the engraftment of hmOEG and hmOEG induced neurons could be a procedure to enhance neural repair after central nervous system injury.

## INTRODUCTION

Since Ramon y Cajal classical studies it is known the scarce regenerative capacity of the Central Nervous System (CNS) (Ramón y Cajal, 1928). Several strategies have been used to overcome this hurdle, such as axonal growth inhibitors blocking agents or transplantation of different cell types (Assinck et al., 2017; Fischer et al., 2020; Martins-Macedo et al., 2020). A recent approach to foster CNS regeneration after injury or disease is to generate new neurons in the damaged site; in this regard, one technique used has been to reprogram somatic cells into pluripotent stem cells (iPSCs) and differentiate them into neurons for cell replacement therapy. However, iPSCs technology has suffered from some drawbacks as demonstrated by tumorigenesis, indeterminate differentiation or genomic instability (Attwood & Edel, 2019; Bellin et al., 2012). An alternative approach to bypass this downside is cellular direct conversion: to reprogram somatic cells into terminally differentiated cells without going through a stem cell stage (Aydin & Mazzoni, 2019; Bocchi et al., 2022; Vignoles et al., 2019). The induced neurons (iNs) are either generated *in vitro* and subsequently transplanted at the site of the lesion or reprogramming is induced from glial cells located at the affected area (Leaman et al., 2022). Most of glia-to-neuron conversion research has been carried out using virus-mediated ectopic expression of neurogenic transcription factors - NEUROG, NEUROD, ASCL1- on their own or in combination with other factors to enhance cell maturation, efficiency or specification (Bocchi et al., 2022; Masserdotti et al., 2016).

The choice of the cell type to be reprogrammed is a major challenge for the potential applications of iNs. Fibroblasts are an abundant and easily accessible cell type and have been successfully converted into various neuronal subtypes, even though it is a cell lineage distantly related to neurons (Caiazzo et al., 2011; Colasante et al., 2015; Wang et al., 2020). Astrocytes are one of the main cell sources for direct conversion to neurons as they share a common neural origin and they are ubiquitously distributed throughout the CNS. Thus, it has been reported that expression of NEUROG2 or ASCL1 in postnatal mouse astrocytes generates glutamatergic or GABAergic neurons (Heinrich et al., 2010; Masserdotti et al., 2015). In vivo astrocyte to glutamatergic neuron conversion has also been accomplished by expression of a single neural transcription factor, NEUROD1 (Chen et al., 2020; Guo et al., 2014) or striatal astrocytes into GABAergic neurons through ectopic expression of NEUROD1 and DLX2 transcription factors (Wu et al., 2020). Other glial cells such as microglia and NG2 have also been reprogrammed to neurons using the NEUROD1 factor (Matsuda et al., 2019) or combining three factors - ASCL1, LMX1, NURR1 - (Pereira et al., 2017; Torper et al., 2015), respectively. Nevertheless, these glial cell types are difficult to obtain from patients.

In this work we selected olfactory ensheathing glia (OEG) as a candidate for direct conversion to neurons. An advantage of OEG over other cell types is the reported capacity of OEG in promoting CNS regeneration (reviewed in Gómez et al.,2018). OEG is located in the mammalian olfactory system and provides a pro-regenerative environment for olfactory sensory neuronal axons (Gómez et al., 2018). In adult lifetime, olfactory sensory neurons are constantly renewed and OEG is responsible for facilitating axonal growth from the neuroepithelium to mitral and tufted cells in the olfactory bulb (Gómez et al., 2018; Roet & Verhaagen, 2014). OEG neuroregenerative capacity, both from olfactory bulb and mucosa, has been tested in an *in vitro* model of axotomized rat retinal ganglion neurons (García-Escudero et al., 2012; Moreno-Flores et al., 2003; Portela-Lomba et al., 2020) and *in vivo* in rat models of spinal cord injury (SCI) (Khankan et al., 2016; Moreno-Flores et al., 2006; Richter et al., 2005; Thornton et al., 2018). Its reparative ability is due to a combination of several factors: these cells express membrane bound and secreted adhesion molecules that promote axonal growth (L1, E-NCAM, laminin, fibronectin) (Gómez et al., 2018); they secrete proteases (MMP2, MMP9, PAI-1) (Pastrana et al., 2006; Simón et al., 2011), being able to reorganize the reactive glial environment (Ramer et al., 2004) and to decrease reactivity and size of the glial scar (García-Alías et al., 2004; Lakatos et al., 2003; Ramer et al., 2004); also, they secrete neurotrophic factors (NGF, BDNF, GDNF) (Pastrana et al., 2007; Woodhall et al., 2001) and cytokines (interleukin 6, TGFβ3) (Roet & Verhaagen, 2014) that play a relevant role in neuroprotection and repair in the damaged zone. Furthermore, these cells can easily be obtained from the olfactory mucosa of patients by biopsy (García-Escudero et al., 2012) so autologous therapies can be performed and avoid post-transplant rejections.

Thus, if we add to the neuroregenerative capacity of OEG cultures the conversion to neurons of a fraction of their population through reprogramming techniques, the engraftment of OEG and OEG induced neurons (OEG-iNs) could enhance neural repair at the damaged site. Previously, Sun et al. have demonstrated that OEG from adult mice can be directly reprogrammed into neuronal cells by the transcription factor NEUROG2 (Sun et al., 2019). Based on these promising studies in rodents, our aim was to generate iNs from human adult OEG. In the present work we characterized a primary OEG culture from adult human olfactory mucosa (hmOEG) that showed neuroregenerative properties. After transduction of the single neurogenic transcription factor NEUROD1, the cells under study not only exhibited morphological and immunolabeling neuronal features but were able to fire action potentials and expressed glutamatergic and GABAergic markers. In addition, after engraftment of transduced hmOEG cells in mouse hippocampus, these cells showed specific neuronal labeling. These findings lay the groundwork for the potential use of hmOEG-iNs in cell-based therapy or disease modeling.

## METHODS

### Animals

Adult male Wistar rats (RccHan®:WIST) were obtained from Envigo (Envigo RMS Spain, SL). Wild type C57BL6 mice were obtained from the animal facility of Universidad Francisco de Vitoria. Immunosupressed NOD-SCID mice were acquired and maintained in the Insitituto de Investigaciones Biomedicas “Alberto Sols” (IIBM) animal facility under sterile conditions.

Animal procedures were performed in animal facilities of Universidad Francisco de Vitoria and Facultad de Medicina of Universidad Autónoma de Madrid. The surgeries of the NOD-SCID mice were carried out in the Instituto de Investigaciones Biomédicas (IIBm-CSIC-UAM) facilities under a flow laminar hood.

All animal procedures were carried out complying with the European Council Directive 2010/63/UE and Spain RD 53/2013, and approved by national and institutional bioethics committees with the authorization code PROEX 142.2/20. Animals were housed under a 12-hour light/12-hour dark cycle and were supplied with ad libitum access to regular food and water.

### hOEG cell cultures

Primary culture of human olfactory mucosa (hmOEG) (García-Escudero et al., 2012) and immortalized human olfactory bulb OEG (ihOEG) cell lines Ts14 (García-Escudero et al., 2011; Lim et al., 2010) and Ts12 (Plaza et al., 2016), were maintained in ME media, composed by DMEM/F12 (Gibco) supplemented with 10 % FBS (GE Healthcare Hyclone), 2 mM glutamine (Lonza), 20 µg/mL pituitary extract (Gibco), 2 μM forskolin (FSK; Sigma) and antibiotics (P/S/A, penicilin/estreptomicin/anfotericin; Lonza) at 37°C in 5% CO2.

### Mouse neuron primary culture

Mouse embryonic neurons were obtained from cerebral cortex of E17 C57BL6 mice using the Worthington Biochemical Corporation papain dissociation kit (ref. LK003150) After removing the meninges, the cortex was transferred to a papain plate, where they were cut in <1 mm pieces with a scalpel blade. Then, they were incubated at 37°C for 30 minutes with intermittent shaking. Afterwards, they were mechanically disaggregated by pipetting with a glass Pasteur 10-15 times until a homogeneous suspension was obtained. Subsequently, in order to remove the remained aggregates, the suspension was passed through a 0,75 mm mesh and then centrifuged for 5 minutes at 200 x g. Supernatant was aspirated and cells were resuspended in 5 mL of NB-B27 Plus medium (Neurobasal Plus (NB, Gibco) supplemented with B27 Plus (Gibco), 2 mM glutamine (Lonza) and P/S/A (Lonza)) and 100.000 cells were plated on glass coverslips (12mm, ThermoScientific) pre-treated with PLL-Laminin (10 µg/mL-5 µg/mL).

### Mouse astrocyte primary culture

Astrocytes were isolated from the cortex of postnatal 0-2 days old (P0-2) C57BL6 mice. Cerebral cortex was dissected and, after removal of meninges, cut with a scalpel blade until <1 mm pieces were obtained. Then, they were passed through a 5 mL pipette 5 times and incubated at 37°C for 30 minutes with intermittent shaking. Afterwards, they were mechanically disaggregated by pipetting with a glass Pasteur 20-25 times until a homogeneous suspension was obtained. Subsequently, the suspension was passed through a 0,75 mm mesh to remove clumps and centrifuged for 5 minutes at 200 x g. Thereafter, the supernatant was aspirated, cells were resuspended in the corresponding volume of M10 medium (1 mL per 4 hemispheres) composed of DMEM/F12 (Gibco) supplemented with 10 % FBS (GE Healthcare Hyclone), 2 mM glutamine (Lonza) and P/S/A (Lonza), and 1 mL of the supernatant was plated in a T75 culture flask (hereafter T75 flask; Falcon), pre-treated with poly-L-lysine (PLL, 10 µg/mL; Sigma). When the cells reached 90-95% confluence, around day 7-10 post-cultured, the T75 flasks were shaken O/N at 37°C to obtain a purified astrocyte culture.

### hOEG co-cultures with retinal ganglion neurons (RGNs)

Cocultures of hOEG with axotomized RGNs were carried out to assess the neurogenerative capacity of hOEG as previously described (Portela-Lomba et al., 2020). Briefly, hOEG cells were plated on 12 mm coverslips in a 24-well plate (M24; Cultek), aiming to have a monolayer the following day. RGNs were axotomized and extracted from retinas of 2 month old adult rats (Wistar rats; Envigo) by sectioning the optic nerves and were dissociated using the papain kit from Worthington Biochemical Corporation. Isolated RGNs were plated on top of the hOEG monolayers and after 96 hours in coculture, these were fixed with 4% paraformaldehyde (PFA) for immunostaining.

To evaluate the regenerative capacity of hOEG over the axotomized RGNs, the cocultures were immunolabeled with SMI31 (MAP1B and neurofilament H axonal markers) and MAP2A&B (somato-dendritic marker) and analyzed with the 40X objective of an inverted fluorescent microscope (DMi8, Leica). Immunofluorescence images were quantified with ImageJ software (ImageJ; NIH) and axon length was measured using the NeuronJ plugin. Quantification of axonal regeneration was determined by calculating: 1) the percentage of neurons with axons, detected with SMI31, versus the total number of neurons, labelled with MAP2A&B; 2) the mean axonal length per neuron, by calculating the ratio of the sum of the lengths (µm) of all axons out of the total number of neurons counted. At least 30 fields were randomly imaged and at least 200 neurons were quantified in each preparation. Experiments were repeated independently at least three times.

### Lentivirus production

The packaging plasmids pMD2.G (#12259) and PAX2 (#12260) were acquired from Addgene. The plasmids pCAG-*Gfp*-IRES-*Gfp* (LV-*Gfp*) and pCAG-*NeuroD1*-IRES-*Gfp* (LV-*NeuroD1*) were obtained in retroviral vectors, kindly donated by Dr. Gong Chen (Guo et al., 2014) and then they were cloned into pRRL.sin.cPPT.CMV.Wpre (Follenzi et al., 2000). pCAG-*Ngn2*-IRES-GFP (LV-*Ngn2*) (Herrero-Navarro et al., 2021) was kindly supplied by Dr. Guillermina López-Bendito. *Ascl1* was cloned into pLenti CMV/TO Neo empty (w215-1) plasmid (Addgene #17485).

For the lentiviral production, 5×10^6^ HEK293T cells were plated in 10-cm plates (p100, Falcon). The following day, cotransfection of 5 µg of pMD2G, 6 µg of PAX2 and 10 µg of the lentiviral vector were carried out using the calcium phosphate method (Kingston et al., 1999) in Optimem media (Gibco). After 6h post transfection, a 10% glicerol shock was performed. First, a solution composed by glicerol:HBS:ddH_2_O (1:5:4) was prepared. Then, the media was aspirated and 2 mL of the glicerol solution was added to each p100 during exactly 2 min. Subsequently, the solucion was aspirated and washed with PBS 1X before adding 7 mL of M10 media per plate. After 72 h post transfection, the media containing the lentiviral particles was harvested, filtered through a low binding 0.45 µm mesh and concentrated at 10 °C in a ultracentrifuge (OPTIMA XE-90) for 2 h at 82.700 g. Finally, the pellet containing the virus was resuspended in PBS 1X and stored at −80 °C. Viral stocks were titered by flow cytometry, after infection of HEK293T cells, to get the % of GFP+ cells. Titer average was 10^7^ transducing units (TU) /ml.

### hmOEG reprogramming

For neuronal reprogramming, 20.000 cells/well of hmOEG were plated in M24 cell plates (M24-I, Ibidi) pre-treated with poly-L-ornitine/laminin (PO/L: 20µg/mL/5µg/mL) to reach 80-90% of confluence. Next day, cells were infected with the lentiviral particles in M10 media containing polybrene (8 µg/mL, Sigma) with a MOI of 10. After 6-20h, the viral media was changed with fresh M10 medium. The following day, the media was replaced with neuronal diferentiation media 1 (NDM1: DMEM/F12 (Gibco), glutamax (Gibco), P/S/A (Lonza), glucose 3,5 mM (Sigma), FBS 1% (Gibco), N2 (Gibco) and B27 (Gibco)) with 10 μM FSK. Media was partially renewed every 3-4 days with neuronal differentiation media 2 (NDM2: 1:1 DMEM/F12:NB (Gibco), glutamax (Gibco), P/S/A (Lonza), glucose 3,5 mM (Sigma), N2 (Gibco), B27 (Gibco) and 20 ng/mL of the maturation factors BDNF, GDNF and NT3) containing 10 μM FSK.

### hmOEG induced neurons (hmOEG-iNs) coculture with astrocytes

Infected hmOEG were cultured onto a monolayer of mouse postnatal astrocytes to increase hmOEG-iNs viability. For the coculture, 75.000 astrocytes were plated in M24-I pretreated with polyornitine (20µg/mL)–laminin (5µg/mL). The remaining astrocytes were plated in 150 mm plates (P150), pretreated with PLL to obtain conditioned NDM2 from the astrocytes. After 48 hours, 20.000 hmOEG-iNs (48h post-infection) were plated onto the astrocytes in ME media. Next day, the medium was replaced by conditioned NDM1 with 10 μM FSK. Every 3-4 days the media was refreshed by conditioned NDM2 containing 10 μM FSK. The cocultures were maintained up to 90 days.

### Cell transplantation

NOD/SCID mice 1 month old were anesthetized with isoflurane. Transduced hmOEG (7 days post-infection) were stereotactically transplanted at a concentration of 200.000cells/2 µL (infusion rate of 0.5 µL/min), into the hippocampus (right hemisphere) in the following coordinates from Bregma: AP-2, ML1.5/-1.5, DV-2. Mice were sacrificed 2- or 3-months post injection for immunohistological analysis.

### Immunostaining

Cell cultures were fixed with 4% PFA, permeabilized and blocked with PBS-TS (PBS, 0.1% Triton, 5% FBS) for 30 minutes at RT and then incubated at 4 °C O/N with the corresponding primary antibody diluted in PBS-TS. Next day, three 5-minute washes were performed with 1X PBS and incubated with the desired fluorescent secondary antibody (Alexa), diluted in PBS-TS, for 1 hour at RT in the dark. Afterwards, they were washed 3 times with 1X PBS for 5 minutes and incubated with DAPI (4’6-Diamidino-2-phenylindole; 1 µg/mL, MERCK) for 5 minutes. After washing, samples were mounted with fluoromount (Southern Biotech) and observed under an inverted fluorescent microscope (DMi8, Leica). Antibodies and antibodies dilutions are listed in Supplemental Table 1. Fields were randomly photographed, and the intensity density of the somas was quantified in an automated and blind way, taking as region of interest the DAPI staining. Immunofluorescence signals were quantified using ImageJ software (ImageJ; NIH).

For the cerebral tissues, mice were anesthetized with 0.2 mL of pentobarbital (33 mg/mL) and then intracardially perfused with 4% PFA. Subsequently, brains were extracted, embedded in *Tissue-Tek OCT Compound*, and sliced in the cryostat at 25 µm. For the immunohistochemistry, the brain slices were blocked with PBS-TS2 (PBS, 0.5% Triton, 10% serum and 1% BSA) for 1h at RT and then incubated with the desired primary antibody at 4 °C O/N. Next day, the brain slices were washed 3 times with PBS for 10 min before the incubation with the fluorescent secondary antibody (Alexa) for 2 h at RT in the dark. Then, three 10-minute washes were performed with PBS, following an incubation with DAPI for 10 min. After washing, brain slices were placed in slides, dehydrated and mounted with DEPEX (MERCK). Samples were examined with a Leica TCS SP5 spectral confocal microscope and for image acquisition, a LAS-AF software was used. Images were taking with 20x and 40x objectives and using 405 nm (DAPI staining) 488 nm (green immunostaining) and 561 nm (red immunostaining) laser lines.

### Electrophysiology

hmOEG-iNs 30-, 60- or 90-days post-infection were recorded using the patch-clamp technique in their whole-cell configuration in voltage-clamp and current-clamp modes. Recordings were made with borosilicate pipettes (OD-ID: 1.5-0.86; Sutter Instrument CO, Novato, CA), with a resistance of 4-8 MΩ and were filled with an intracellular solution (5 mM KCl, 20 mM HEPES, 2 mM CaCl2, 2 mM MgCl2, 0.6 mM EGTA, 130 mM K-gluconate, 2 mM ATP, 0.2 mM GTP, 20 mM phosphocreatine and 50 U/mL creatine phosphokinase; pH 7.3 and osmolarity 286 mOsm). Recordings were performed at a temperature of 30°C with a constant flow of 2mL/minute extracellular solution (150 mM NaCl, 2.5 mM KCl, 10 mM HEPES, 30 mM glucose, 1 mM MgCl2 and 2 mM CaCl2; pH 7.3). Action potentials and sodium currents were evoked by depolarizing pulses of current and voltage. Cells were accepted only when the seal resistance was above 1 GΩ and the series resistance did not change by 20 % during the experiment. Signals were filtered at 3 KHz and sampled at 10 KHz with a Digidata 1500A analogue-to-digital conversion card (Molecular Devices, Sunnyvale, CA). Mice embryonic cortex neurons cultured *in vitro* between 7-14 days were recorded as a positive control. After obtaining a stable recording of action potentials and sodium currents, recordings were blocked by adding tetrodotoxin (TTX; 0.5 µM, Abcam), a blocker of voltage-dependent sodium channels, to the extracellular solution.

### Statistical analysis

For statistical comparisons between two populations Student t test was applied. For multiple comparisons, one way-ANOVA and post hoc Tukey test was applied. To analyze the intensity density of the immunofluorescences, two-factor analysis of variance (two-way ANOVA) followed by the multiple comparison between means Sidak’s post-hoc test was applied. Statistical significance was established at a value of p<0.05.

## RESULTS

### 1. Primary cultures from adult human olfactory ensheathing glia show neuroregenerative properties

OEG primary cultures were prepared from olfactory mucosa from a thirty-year-old donor (García-Escudero et al., 2012), hereafter hmOEG. To verify the identity of this cell line we carried out an expression analysis of OEG markers - S100 glial calcium binding protein B (S100B), glial fibrillary acid protein (GFAP) and vimentin - by immunofluorescence techniques. As a positive control we used an olfactory bulb immortalized human OEG cell line (TS14, from now on) previously described in (García-Escudero et al., 2011; Plaza et al., 2016). Almost 100% of the hmOEG cells expressed the cytoplasmic markers S100B (Figure S1: A’, B’, C) and vimentin (Figure S1: A’’’, B’’’, C). In addition, 62±13% of hmOEG and 67±20% of TS14 cells were GFAP positive although immunoreactivity against GFAP was rather diffuse in both cell types (Figure S1: A’’, B’’, C).

We then verified that hmOEG cells did not express neuronal markers by labeling β3-tubulin (TUBB3) (Tuj1) and neuronal nuclear antigen NeuN/FOX3. Mouse embryonic cortex neurons were used as a positive control. Expression of NeuN was absent in the hmOEG line (Figure S2: ’’’, B’’, C) and 25% of the population stained positive for Tuj1 but with a more diffuse pattern compared to the positive control (Figure S2: ’’, B’, C). To rule out the presence of neuronal precursors we performed immunofluorescence to detect SOX2, expressed in neural progenitor cells. As a positive control we used the above mentioned neuronal primary culture from mouse embryonic cortex, which due to its embryonic stage is a niche for neuronal precursors. Our results excluded the presence of neuronal precursors in our hmOEG cell line: the embryonic cortex culture contained 19±6% of SOX2-positive neuronal precursors (Figure S2: ’’’, C) but hmOEG cells did not express SOX2 (Figure S2: B’’’, C).

Next, we assessed the regenerative properties of hmOEG. For this purpose, we conducted an *in vitro* regeneration assay in a co-culture of hmOEG with axotomized adult rat retinal ganglion neurons (RGNs) (see Methods section). TS14, a line with regenerative capacity, and TS12, an immortalized OEG line with low regenerative capacity, were used as controls (Plaza et al., 2016); as a negative control we grew axotomized RGN cells on a poly-L-lysine (PLL) substrate to verify that they did not inherently extend axons. Axonal regeneration was quantified using immunostaining against SMI31 (a phosphoepitope of neurofilament H and MAP 1B) as an axonal marker and against MAP2A/B to identify the somatodendritic compartment. Two parameters were assessed to determine regenerative capacity: the percentage of RGNs that extended an axon and the mean axonal length/neuron of these axons. We observed that the regenerative capacity of hmOEG was similar to our positive control: percentage of neurons with an axon was 14±1% and the average axonal length was 38±4 micrometers/neuron (Figure 1). Furthermore, a statistically significant higher regenerative capacity of both parameters was observed with respect to the low regenerative Ts12 and negative PLL controls (Figure 1).

**FIGURE 1.**
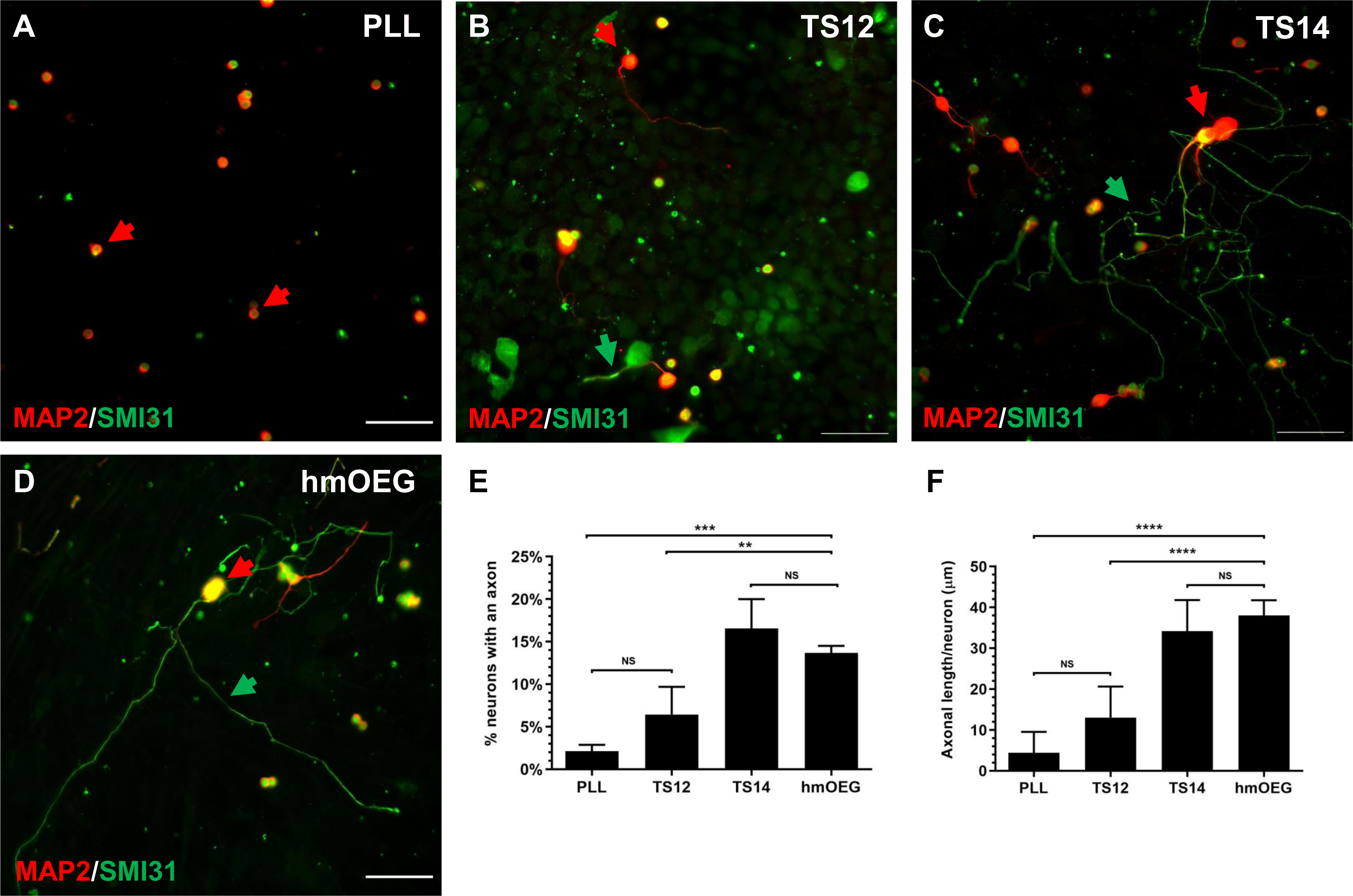
hmOEG shows regenerative capacity in an *in vitro* model of adult axonal regeneration. (A–D) show representative images of axotomized retinal ganglion neurons after 96 h in co-culture with OEG. OEG human cell line TS14 was used as a positive control for neuro-regeneration (C) and human cell line TS12 (B) and PLL (A) correspond to low regeneration and negative controls, respectively; (D) corresponds to hmOEG. Green arrows indicate axons positive for axonal marker SMI31 (green) and red arrows indicate somas and dendrites positive for somatodendritic marker MAP2 (red). Histograms represent the mean ± SD of the quantifications: percentage of retinal neurons extending an axon (E) and the mean axon size expressed as μm/neuron (F). Statistical tests applied were one-way ANOVA and post-hoc Tukey test (****p ≤ 0.0001; ***p ≤ 0.001; NS, not significant) for multiple comparisons between means (n = 4, per experiment ≥ 30 fields were analyzed). Scale bar: 50 μm.

In summary, the hmOEG primary cell line expressed OEG markers but did not express neuronal or neuronal precursor markers and, moreover, showed pro-regenerative properties in an *in vitro* model of adult axonal regeneration.

### 2. Ectopic expression of NEUROD1 directly converts hmOEG to neuronal cells

We selected the transcription factors NEUROG2, NEUROD1 and ASCL1, which are proteins of the bHLH family of transcription factors involved in specification and neural differentiation, for screening to convert hmOEG to induced neurons (hmOEG-iNs). hmOEGs were infected with lentiviral particles to express the candidate factors, individually or in combination with each other (see Methods section) and were identified by their co-expression with the green fluorescent protein (GFP). As a negative control we infected the cells with the same lentivector, but the transcription factor gene was replaced with a second copy of the GFP gene (LV-GFP from now on).

We assessed the reprogramming competence of the transcription factors by immunostaining with the pan neuronal marker Tuj1 in GFP-positive cells, 30 days post-infection (dpi). Ectopic expression of NEUROD1 induced Tuj1 expression and a morphological change towards a neuronal phenotype (Figure S3: B, B’). By contrast, neither LV-GFP nor the other candidate genes triggered conversion of hmOEG into Tuj1-positive neuronal cells (Figure S3: A,A’, C,C’, D,D’); indeed, when we overexpressed ASCL1 alone or in combination with the other factors, cell death occurred in infected cells (data not shown). Based on these results, we selected the transcription factor NEUROD1 to study the reprogramming process.

To determine the reprogramming efficiency of NEUROD1 ectopic expression, we calculated the percentage of Tuj1-positive cells in relation to the total number of infected, GFP-positive cells (Tuj1+/GFP+). At 7 dpi the number of Tuj1 expressing cells constituted 31±5% of the infected cells (Figure 2 A, ’’, I), reaching 58±2% after 30 dpi (Figure 2 D, D’, I). In addition, representative images show a change in cell morphology along time, with a gradual development of neuronal-like extensions (Figure 2 A-D’). Immunostaining of the mature neuronal marker NeuN showed positive signal after 21 dpi (Figure 2 E-G’, J) but it was not until 30 dpi when a significant increase (39±11%) of GFP positive cells expressing NeuN was observed (Figure 2 H, H’, J).

**Figure 2.**
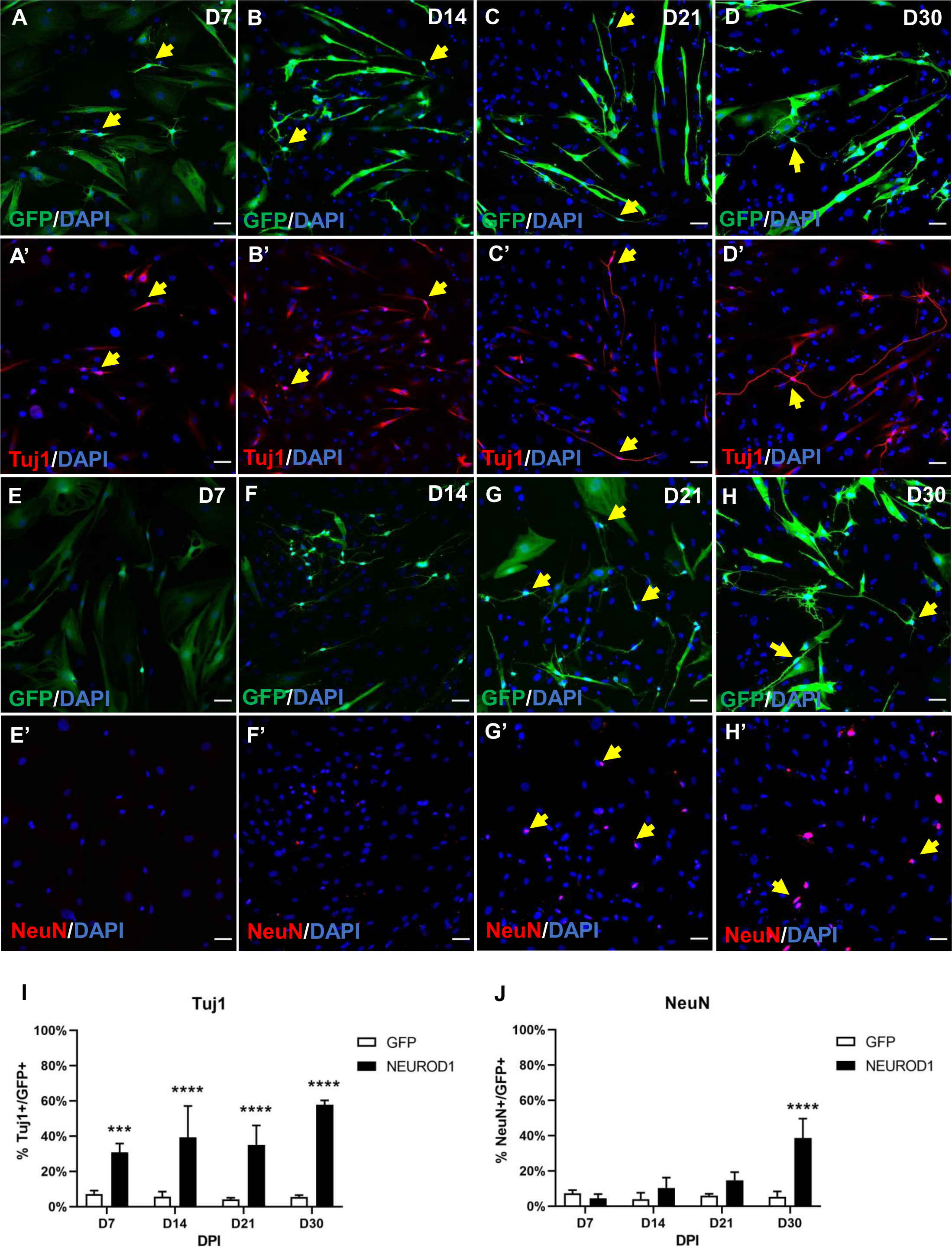
NEUROD1-hmOEG express neuronal markers. Representative images of the neuronal markers Tuj1 (A-D’) and NeuN (E-H’), expressed at 7, 14, 21 and 30 dpi. Note the neuronal morphology at 30dpi. GFP+ cells carry the NEUROD1-GFP construct. Yellow arrows indicate cells expressing the marker. The histograms (I, J) represent the mean±SD of the percentage of Tuj1 or NeuN positive cells in relation to the total number of infected, GFP+ cells (NEUROD1: carry the NEUROD1-GFP construct, GFP: carry the GFP-GFP construct). Statistical tests applied were Two-way ANOVA and post-hoc Sidak test (****p≤0.0001,***p≤0.001) for multiple comparisons between means (n=3 independent experiments, 25 fields were analyzed). Scale bar: 50 µm. dpi: days post-infection.

We further examined neuronal maturation by immunocytochemical analysis of the axonal marker SMI31, whose expression in hmOEG-iNs began at 14 dpi and was held over time (Figure 3 A-D’). It is worth noting that at 30 dpi, hmOEG-iNs expressed SMI31 in a positive gradient from the soma towards the axon ends (Figure 3 D’), a pattern characteristic of mature neurons. Likewise, hmOEG-iNs stained positive for the presynaptic neuronal marker synapsin from 21 dpi onwards, with a characteristic expression in discrete puncta along the axon, at 30 dpi (Figure 3 E-H’).

**Figure 3.**
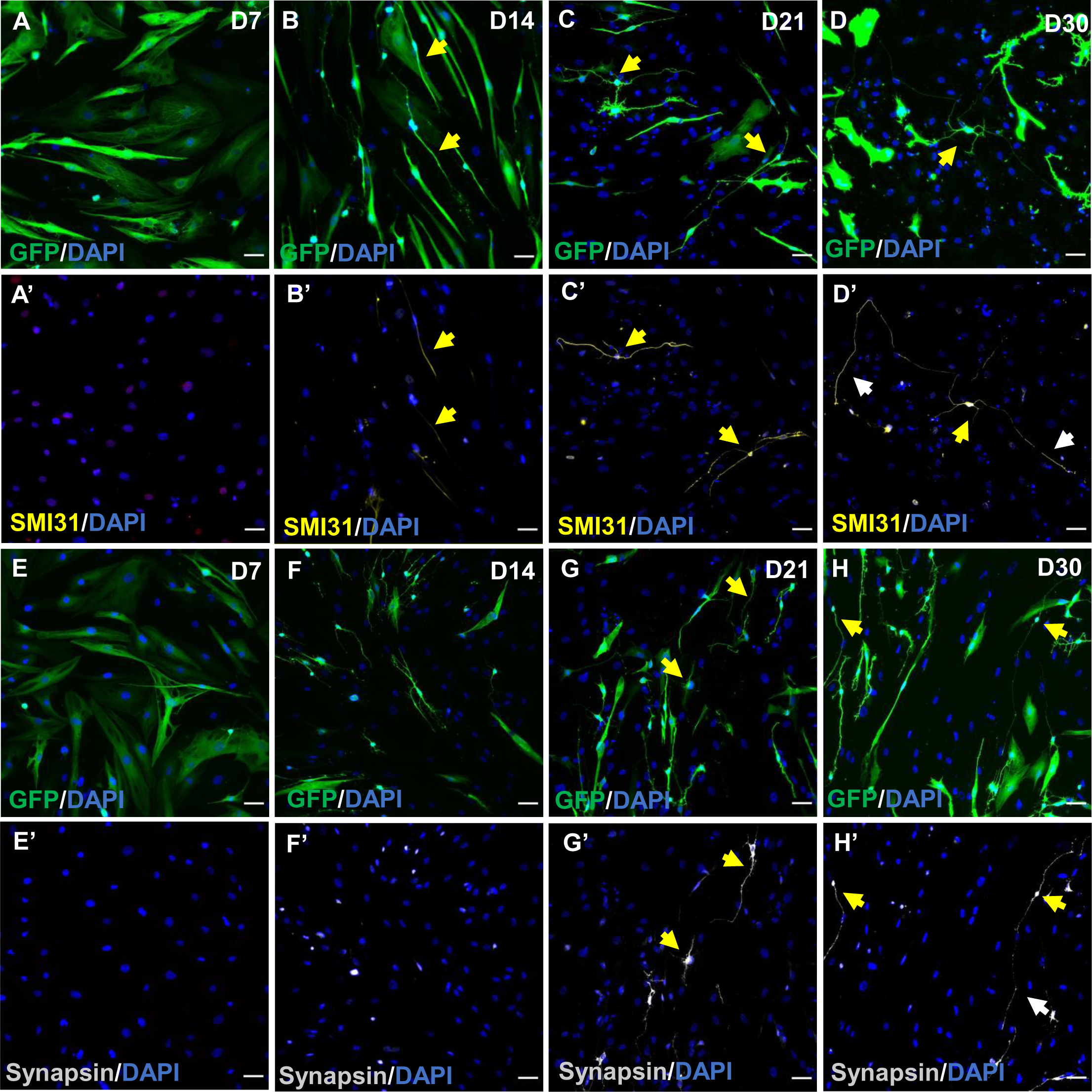
Expression analysis of mature neuronal markers in NEUROD1-hmOEG. Representative images of neuronal markers SMI31 (A-D’) and synapsin (E-H’), expressed at 7, 14, 21 and 30 dpi. Yellow arrows indicate cells expressing the marker. Of note is the pattern of SMI31 expression in a positive gradient from the soma towards the axon ends (D’, white arrows), a characteristic axonal pattern. Synapsin labelling along the axon is also visible (H’, white arrowheads). GFP+ cells carry the NEUROD1-GFP construct. Scale bar: 50 µm.

It has been proposed that cells in the reprogramming trajectory could pass through a neural stem cell-like state, prior to differentiating into iNs (Morris, 2016). This dedifferentiated stage is characterized by transient expression of genes that are normally expressed in neural stem or progenitor cells during embryonic development (Karow et al., 2018; Treutlein et al., 2016). To study whether the reprogramming process of hmOEG to neurons involves a neuroprogenitor intermediate we analyzed the expression of SOX2, a characteristic marker of neuronal precursors, during the first days of induction (2, 4 and 6 dpi). In parallel, we also assessed the expression of the cell proliferation marker KI67.

Immunocytochemical analysis showed that 72±7% % of NEUROD1 infected hmOEG were in a proliferative stage at 2 dpi, but at days 4 and 6 dpi there was a significant drop in the percentage of cells expressing KI67 (Figure S4), suggesting that there was no expansion of progenitor cells during the hmOEG-to-neuron conversion. Regarding SOX2 expression, there was no significant variation over the days in the percentage of positive cells for this neural precursor marker (Figure S4), suggesting that NEUROD1 induced cells do not go through a dedifferentiation stage.

In summary, the infection of hmOEG with LV-*Neurod1* induces the acquisition of a neuronal phenotype over time, as evidenced by a neuronal morphology and the expression of mature neuronal markers, without transitioning through a neuroprogenitor condition.

### 3. Functional maturation of hmOEG-iNs

To enhance the functional maturation of hmOEG-iNs we co-cultured these cells over a monolayer of postnatal mouse astrocytes. After 60 dpi, we analyzed the culture by immunofluorescence and observed several GFP-positive cells acquiring a mature neuron morphology, with long axonal-like extensions, shorter dendritic-like extensions, and an enlargement at the end of some of the projections that could be associated with growth cones (Figure 4A). Under these conditions hmOEG-iNs expressed the neuronal markers Tuj1, synapsin and, at the beginning of the axon, ANK3 (Figure 4B-C’). ANK3 is a specific marker of the axonal initial segment and is required for the normal clustering of voltage-gated sodium channels at the axon hillock and for action potential firing, thereby proving the maturation status of hmOEG-iNs.

**FIGURE 4.**
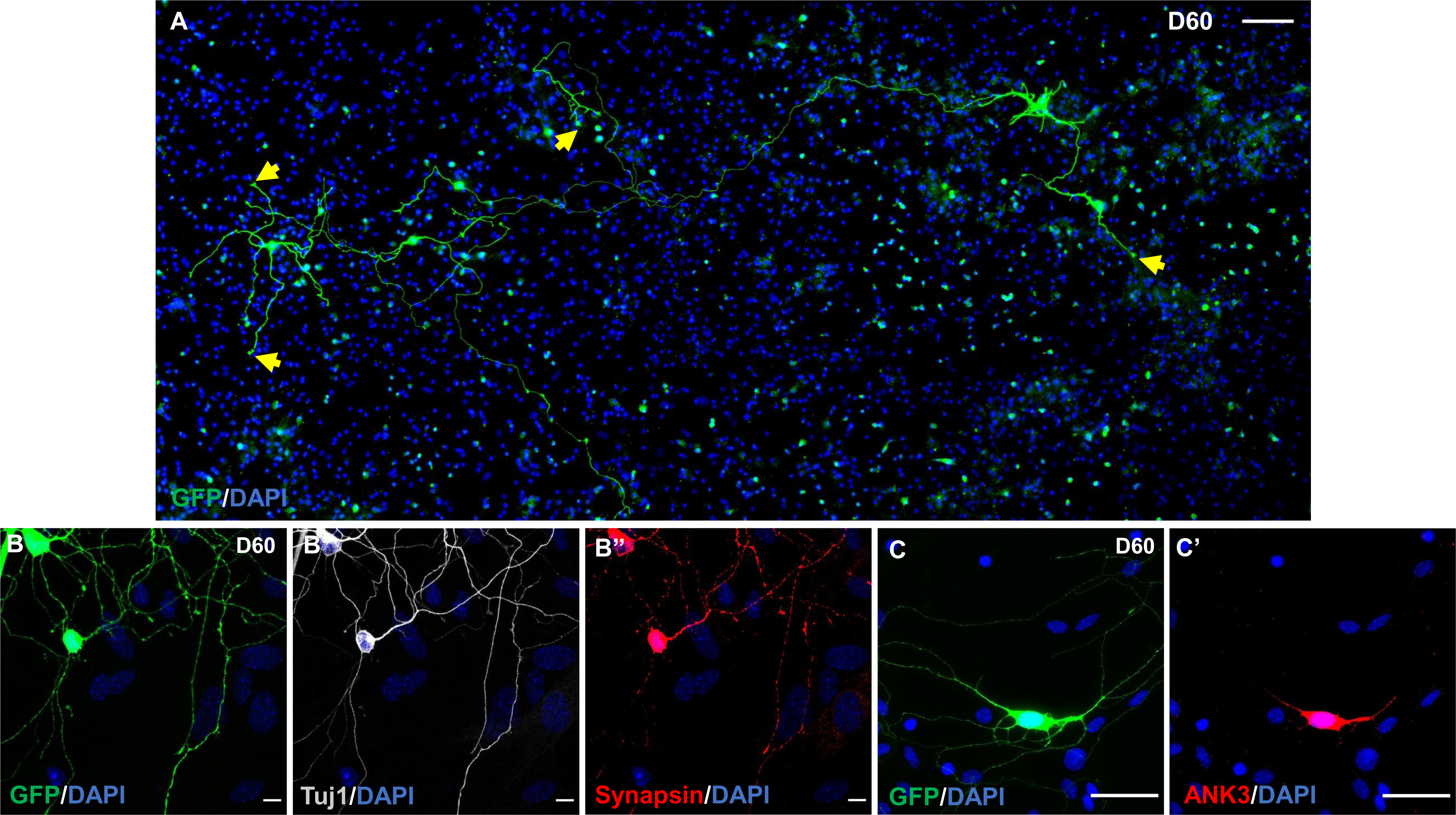
Maturation of hmOEG-iNs. (A) Tilescan of hmOEG-iNs (GFP positive) over a monolayer of cortical astrocytes at 60 days post-infection. Notice the complex neuronal morphology; the yellow arrows highlight growth cone-like thickenings at the end of the extensions. (B-B’’) Under these conditions, hmOEG-iNs (GFP positive) express the neuronal markers Tuj1 and synapsin I. (C-C’) Induced neurons also express ANK3, a specific marker of the axonal initial segment, proving the maturation status of hmOEG-iNs. Scale bar: 100 µm (A), 50 µm (C,C’) and 10 µm (B-B’’).

To determine the functionality of hmOEG-iNs, we performed electrophysiological assays using the patch-clamp technique. Recordings were performed on hmOEG-iNs identified with the GFP reporter, maintained in culture for 60 and 90 dpi. As a positive control we used embryonic mouse cortex neurons (Figure 5A). Recording of hmOEG-iNs 60 dpi revealed a sodium current of 200 pA, sufficient to trigger an action potential. At 90 dpi hmOEG-iNs showed a higher functional competence, recording an increase in sodium current up to 1000 pA and firing a train of action potentials (Figure 5A).

**Figure 5.**
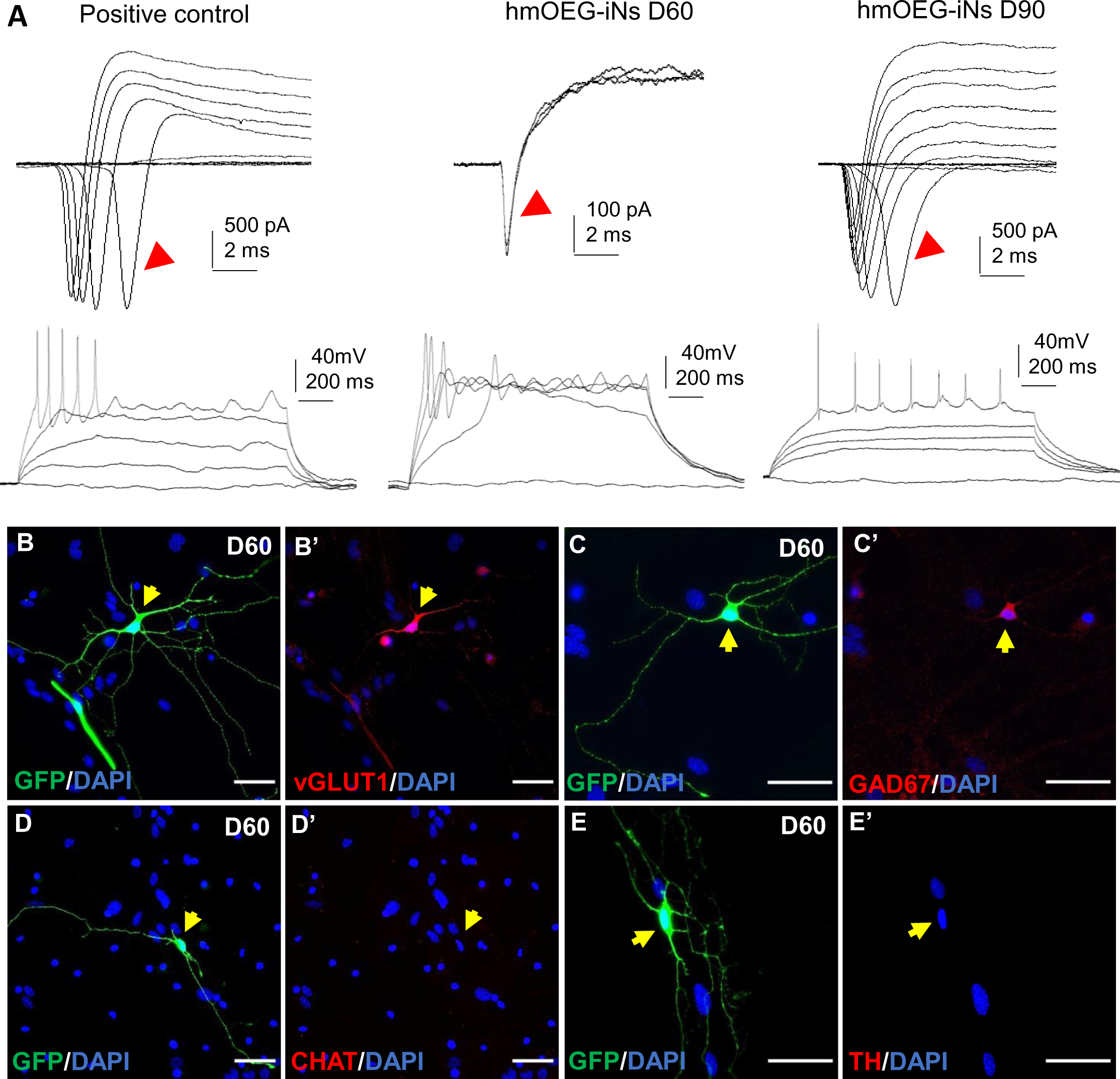
Functional maturation of hmOEG-iNs. (A) Electrophysiological recordings of cortical neurons (left, as positive control), induced neurons at 60 dpi (middle) and induced neurons at 90 dpi (right). Notice that at 90 dpi, hmOEG-iNs have a sodium influx enough to fire a train of action potentials. Red arrows show the sodium channel currents. (B-E’) Immunofluorescence analysis of neuronal subtype markers: hmOEG-iNs expressed the glutamatergic neuron marker vGLUT1 (B, B’) and the GABAergic neuron marker GAD67 (C, C’), however, no cells stained positive for the markers CHAT (D, D’) or TH (E, E’), for cholinergic and dopaminergic neurons, respectively. Yellow arrows show hmOEG-iNs. Scale bar: 50 µm. dpi: days post-infection.

The neuronal identity of hmOEG-iNs was assessed analyzing the expression pattern of different neuronal subtype markers. After 60 dpi, we detected hmOEG-iNs that expressed the glutamatergic neuron marker vGLUT1 (vesicular glutamate transporter 1) (Figure 5B, B’) and GAD67 (glutamate decarboxylase 67) (Figure 5C, C’), a GABAergic neuron marker. However, no cells stained positive for the markers CHAT (choline acetyltransferase) (Figure 5D, D’), a marker specific for cholinergic neurons, or TH (tyrosine hydroxylase) (Figure 5E, E’), a marker for dopaminergic neurons.

To enhance and facilitate neuronal direct reprogramming, several authors have turned to the use of small molecules that target neurogenic signaling pathways (Bocchi et al., 2022; Vasan et al., 2021; Wang et al., 2022). For such purpose, we added different combinations of small molecules to the NEUROD1-transduced cells (summarized in Supplemental Table 2). Also, we tried to reprogram hmOEG with small molecules alone, without overexpression of any transcription factors (Supplemental Table 3). In addition, we tried the small molecule cocktails published by other research groups (see Supplemental Table 4). Of all the combinations we tested, none managed to improve our standard reprogramming conditions with the transcription factor NEUROD1 and the addition of forskolin (see Methods).

Taken together, these data indicate that ectopic expression of the single transcription factor NEUROD1 in hmOEG induces their direct conversion into glutamatergic and GABAergic functionally mature neurons, without addition of small molecules enhancing reprogramming.

### 4. *In vivo* assay of hmOEG reprogramming

After achieving the direct conversion of hmOEG to functional neurons *in vitro*, we wished to verify that the committed reprogramming could take place *in vivo*. To do this, we infected the hmOEG primary cell line with the NEUROD1 factor and, one week after induction it was injected into the hippocampus of immunosuppressed NOD/SCID mice (detailed in Methods); two and three months after transplantation, we analysed the ectopic hmOEG-iNs by immunofluorescence. In the injection site and deeper in the hippocampus we found GFP-NEUROD1-positive cells, of human origin (Stem121 human cell marker positive staining), along the time course (Figure 6A, E). More in detail, some of these cells integrated into the brain tissue and expressed the neuronal markers Tuj1 (Figure 6B, F), MAP2 (Figure 6C, G) and NeuN (Fig. 6D, H), suggesting that they might have been converted into neurons after transplantation. Unexpectedly, although NOD/SCID mice have a high degree of immunosuppression, a fraction of the ectopic hmOEG-NEUROD1 cell population was phagocytosed by microglia, in an environment of intense astroglial reactivity around the injection site (Supplementary Fig. S5 A, B and C). These results show that after NEUROD1 lentiviral induction *in vitro*, hmOEG successfully integrate in brain tissue and neuronal fate commitment is maintained in an *in vivo* environment.

**Figure 6:**
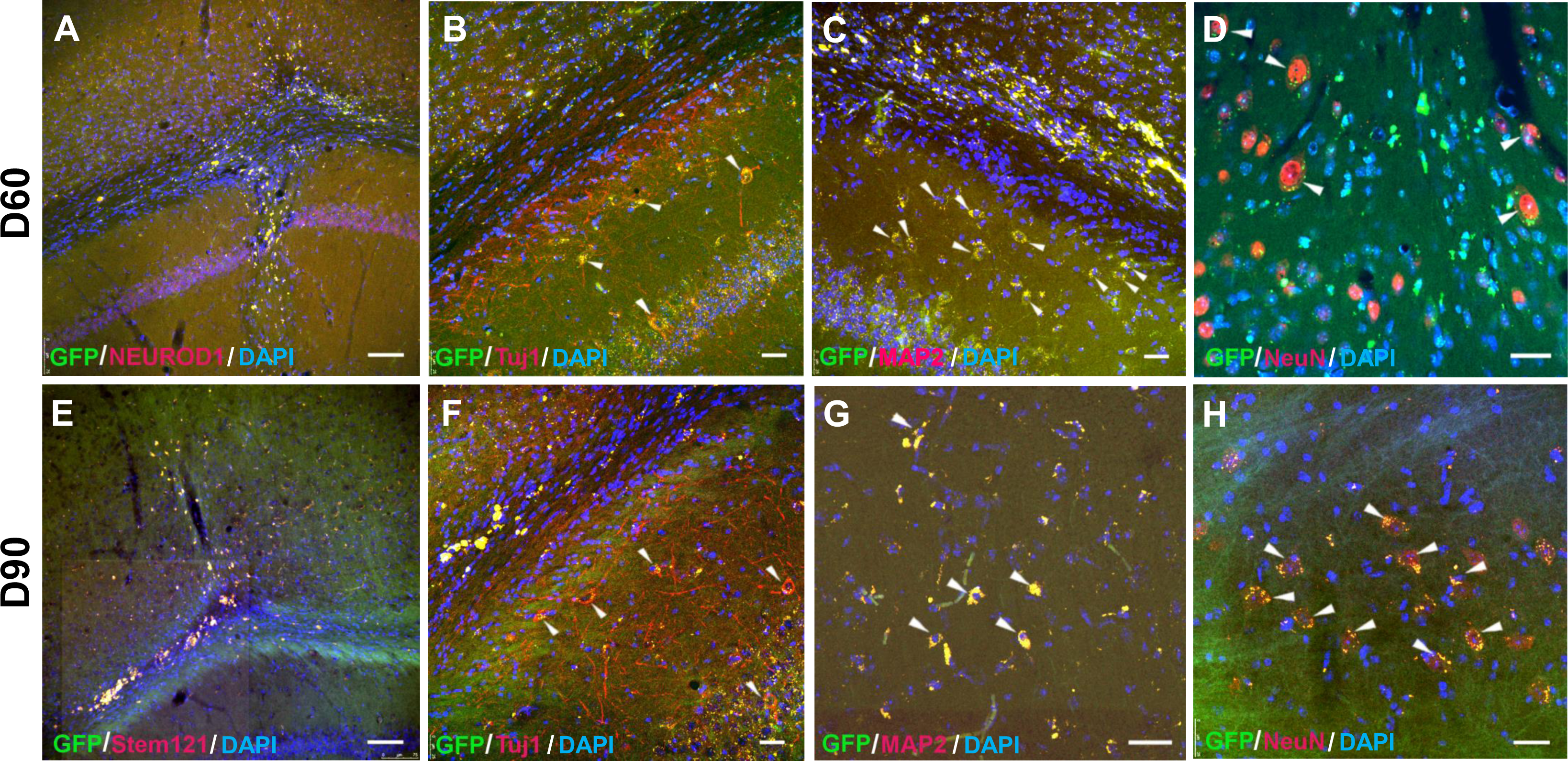
Transplantation of hmOEG-iNs into hippocampus of NOD-SCID mice. (A,E) Survival of hmOEG-iNs at 60 and 90 days post-transplant. GFP+ cells (green) co-express NEUROD1(red) or Stem 121 (red) human antigen, verifying the ectopic origin of these cells. (B-D, F-H) hmOEG-iNs integrate into the brain tissue and stain positive for Tuj1 (red), MAP2 (red) and NeuN (red) neuronal markers (white arrowheads), suggesting conversion into neurons after transplantation. Nuclei were stained with DAPI (blue). Scale bar: 75 µm (A,E) and 25 µm (B-D, F-H).

## DISCUSSION

A recent approach to promote CNS regeneration after injury or disease is direct conversion of somatic cells to neurons, following transplantation to the affected area (Bocchi et al., 2022). In this work we directly reprogrammed olfactory ensheathing glia from adult human olfactory mucosa (hmOEG) to functional neurons, after transduction with the single neurogenic transcription factor NEUROD1, and verified that the committed reprogramming could take place *in vivo* after engraftment of transduced hmOEG cells.

The choice of the cell type to be reprogrammed is a major challenge as it must be easily accessible and susceptible to efficient reprogramming. An advantage of hmOEG over other cell types already reprogrammed to neurons is the reported capacity of OEG in promoting CNS regeneration (Gómez et al., 2018). OEG can be obtained from two sources, from the olfactory bulb and from the olfactory mucosa. No major differences in their neuroregenerative capacity have been demonstrated to date (García-Escudero et al., 2010, 2012), although their genetic profiles have been shown to vary (Guérout et al., 2010) and they promote differential axon sprouting with regard to the axonal tract, after transplantation in the injured spinal cord (Richter et al., 2005)

Therefore, with a view to possible clinical application, we chose hGEO from olfactory mucosa because a biopsy from the olfactory mucosa is less invasive than from the olfactory bulb, so autologous therapies could be performed and avoid post-transplant rejections.

The identity of hmOEG was verified by immunofluorescence analysis of OEG markers and we ruled out the presence of neuronal precursors as well as the expression of neuronal markers in hmOEG prior to their induction into neurons. Unexpectedly, about 30% of hmOEG expressed the general neuronal marker Tuj1, but in a weaker pattern than that of neurons. Previous studies have shown that several OEG lines have a diffuse cytoplasmic expression of Tuj1 (García-Escudero et al., 2012, 2015). Likewise, studies in different cultures of human astrocytes have described that they present a constitutive expression of Tuj1 that is not associated with cell passaging (Dráberová et al., 2008; Knight & Serrano, 2017).

A major feature determining the choice of OEG is its inherent proregenerative capacity (Gómez et al., 2018) and we evaluated this property in co-cultures of hmGEO with axotomized adult rat retinal ganglion neurons (RGNs). In these assays, two parameters were analyzed - the percentage of RGNs extending axons and the mean axonal length/neuron of these axons - whose values were in accordance with previous studies (Plaza et al., 2016; Portela-Lomba et al., 2020).

Most of glia-to-neuron conversion research has been carried out using virus-mediated ectopic expression of neurogenic transcription factors, on their own or in combination with other factors. (Bocchi et al., 2022; Masserdotti et al., 2016). In our work, after transduction of hmOEG cells with lentiviral particles expressing the candidate factors NEUROG2, NEUROD1 and ASCL1, individually or in combination with each other, only ectopic expression of NEUROD1 induced the transition towards a neuronal phenotype. Consistent with our results, previous work shows the effectiveness of NEUROD1 inducing astrocyte to neuron reprogramming (Chen et al., 2020; Guo et al., 2014; Zhang et al., 2020). However, our results are not in agreement with those previously published by Sun et al. (Sun et al., 2019) which demonstrated that OEG from adult mice can be directly reprogrammed into neuronal cells by the transcription factor NEUROG2; moreover, NEUROD1 expression did not result in neuronal conversion. This discordance could be due to a different mechanism inducing cell reprogramming between both species. Thus, RNA expression diversity between humans and mice (Lin et al., 2014), could provide different contexts, in terms of endogenous transcriptional networks and the chromatin state of the starting cell, affecting the reprogramming cell fate (Aydin & Mazzoni, 2019; Ninkovic & Götz, 2018).

A concerning issue was the low reprogramming efficiency after 60-90 dpi, which was accompanied by the death of those cells that did not make it to the iN fate. This effect could be attributed to the metabolic shift during the conversion process, increasing the production of reactive oxygen species (ROS) and inducing cell death by oxidative stress (Gascón et al., 2016). Another process suggested to hinder direct reprogramming is genomic stress, which is caused by interference between high levels of transcription and replication, resulting in high cell death (Babos et al., 2019). Therefore, induction will occur in most cells, but few cells would be fully reprogrammed. Additionally, the starting cell type conversion efficiency will depend on transcriptional accessibility of target genes in a permissive epigenetic environment. Therefore, co-factors that remove epigenetic barriers to reprogramming (e.g. repressive DNA methylation and histone modifications) can enable a neuronal gene expression program and improve transcription factor-mediated neuronal direct conversion (Leaman et al., 2022).

Under appropriate culture conditions, neuronal maturation was achieved, obtaining electrophysiologically competent neurons which expressed glutamatergic or GABAergic markers. It is unclear whether a specific neuronal subtype can be preferentially induced from direct reprogramming of heterologous cells. In previous works, overexpression of NEUROD1 resulted in glutamatergic or, to a lesser extent, GABAergic neurons, starting from astrocytes, NG2 glia or microglia (Guo et al., 2014; Matsuda et al., 2019). Likewise, induction of mouse OEG reprogramming using NEUROG2 resulted in a mixed population of glutamatergic and GABAergic neurons (Sun et al., 2019). In addition, it has been possible to direct cell reprogramming towards a particular neuronal subtype by adding transcription factors involved in cell specification during neuronal development. Thus, reprogramming of striatal astrocytes to GABAergic neurons was induced by a combination of NEUROD1 and DLX2 (Wu et al., 2020).

A reported feature of direct conversion is the lack of stemness of the induced somatic cell population along the reprogramming process. Thus, functional neurons from fibroblasts, astrocytes or microglia have been generated without reverting to a progenitor cell stage (Colasante et al., 2015; Masserdotti et al., 2015; Matsuda et al., 2019). However, it has been proposed that cells in the reprogramming trajectory could pass through a neural stem cell-like state, prior to differentiating into iNs (Cates et al., 2021; Karow et al., 2018; Treutlein et al., 2016). This dedifferentiated stage is characterized by transient expression of genes that are normally expressed in neural stem or progenitor cells during embryonic development. After the analysis of SOX2 expression, a characteristic marker of neuronal precursors during the first days of induction, and of the cell proliferation marker KI67, our results suggested that the reprogramming process did not go through a proliferative dedifferentiation stage. Such intermediate, stem cell-like state was not detected either in murine OEG to neuron direct conversion (Sun et al., 2019).

To enhance neuronal direct reprogramming, several authors made use of small molecules that target neurogenic signaling pathways (Ma et al., 2021; Ninomiya et al., 2023; Wang et al., 2022). This approach overcomes the concern of introducing foreign genetic material to cell cultures that might be potentially used for cell-based therapy. We added different combinations of small molecules to the NEUROD1-transduced cells or to hmOEG without overexpression of any transcription factors, but no combination improved the reprogramming results obtained with the transcription factor NEUROD1. This scenario has already been addressed by some authors (Vasan et al., 2021; Wang et al., 2022) who note the challenge to seek an optimal formula to achieve acceptable conversion efficiency: not only the optimal concentration of each small molecule in the cocktails is difficult to determine but also the length of the induction time is a concern, as longer times can have toxic effects on the cells, while shorter times might not be effective in inducing neuronal conversion.

Induced neurons (iNs) generated in vitro can be subsequently transplanted at the site of the lesion (Leaman et al., 2022). After achieving the direct conversion of hmOEG to functional neurons *in vitro*, we engrafted transduced hmOEG cells in mouse hippocampus: these cells showed specific neuronal labeling, suggesting their conversion to neuronal cells after transplantation. However, a fraction of the ectopic hmOEG-NEUROD1 cell population was phagocytosed by microglia, in an environment of intense astroglial reactivity around the injection site, despite the use of NOD-SCID immunosuppressed mice. Although unexpected, rejection of autologous iPSCs grafts has already been reported (Zhao et al., 2011) and this immune response has been suggested to be triggered by epigenetic alterations present in the reprogramming cell lines (Liu et al., 2017).

In conclusion, in this work we succeeded in directly converting olfactory ensheathing glia from adult human olfactory mucosa (hmOEG) to neurons. We showed that the cells under study maintained the characteristic neuroregenerative properties of OEG and, after transduction of the single neurogenic transcription factor NEUROD1, they exhibited morphological and immunolabeling neuronal features, fired action potentials and expressed glutamatergic and GABAergic markers. In addition, after engraftment of transduced hmOEG cells in mouse hippocampus, these cells showed specific neuronal labeling. Thereby, if we add to the neuroregenerative capacity of OEG cultures the conversion to neurons of a fraction of their population through reprogramming techniques, the engraftment of OEG and OEG induced neurons (OEG-iNs) could be a procedure to enhance neural repair after central nervous system injury.

## ACKNOWLEDGEMENTS

Authors want specially to thank Profs. ’’ngel Nuñez, Francisco Wandosell and Jesús ’’vila for its generous contribution to the financial support of the experiments and for their comments on our work. We thank M. Dolores Morales García (SIdI-UAM Confocal Microscopy Laboratory; Madrid), for her excellent technical assistance in the acquisition of the confocal microscope images, and to Esther García for her excellent technical help in viral production.

## ETHICS APPROVAL STATEMENT

All animal procedures were carried out complying with the European Council Directive 2010/63/UE and Spain RD 53/2013, and approved by national and institutional bioethics committees (Facultad de Medicina, Universidad Autónoma de Madrid and Facultad de Ciencias Experimentales, Universidad Francisco de Vitoria) with the authorization code PROEX 142.2/20.

## AUTHOR CONTRIBUTIONS

MP-L performed experiments and data analysis. DS contributed to the performance of experiments and data analysis. MC-M and GdlF provided excellent technical assistance with the electrophysiology, the *in vivo* studies and the immunohistochemistry, respectively. DF-S performed the electrophysiological experiments and analyses. VG-E contributed to the original project conception, experimental design and performance and to data analysis. JS and MTM-F are main responsible for the conception, funding, experimental design, data interpretation and supervision of the project. JS wrote the first draft of the manuscript. All authors reviewed the article critically for important intellectual content. All authors approved the submitted version.

