## Supplemental Tables for "GENERATION OF FUNCTIONAL NEURONS FROM ADULT HUMAN MUCOSAL OLFACTORY ENSHEATHING GLIA BY DIRECT LINEAGE CONVERSION"

| Table 1. Antibodies | | | | | |
|---|---|---|---|---|---|
| **ANTIBODIES** | **SOURCE** | **COMPANY** | **REFERENCE** | **DILUTION** | |
| | | | | ***IN VITRO*** | ***IN VIVO*** |
| **Ankirin 3 (Ankyrin G)** | MC-Mouse | Thermo Fisher | 33-8800 | 1:250 | - |
| **ChAT** | PC-Goat | Merk | AB144P | 1:1000 | - |
| **GAD67** | MC-Mouse | Millipore | MAB5406 | 1:1000 | - |
| **GFAP** | PC-Guinea Pig | Synaptic Systems | 173002 | 1:1000 | 1:500 |
| **GFP** | PC-Rabbit | Life Technologies | A11122 | 1:1000 | 1:1000 |
| **GFP** | MC-Mouse | Abcam | ab1218 | 1:500 | 1:500 |
| **GFP** | PC-Chicken | Abcam | ab13970 | 1:1000 | - |
| **hNu** | MC-Mouse | Chemicon/Millipore | MAB1281 | - | 1:400 |
| **Iba-1** | PC-Rabbit | FUJIFILM Wako | 019-19741 | - | 1:500 |
| **Ki67** | MC-Mouse | Agilent | M7240 | 1:100 | - |
| **MAP2A&B** | PC-Rabbit | From Dr. Ávila´s Lab | - | 1:500 | - |
| **MAP2** | PC-Chicken | Abcam | ab5392 | - | 1:1000 |
| **NeuN** | MC-Rabbit | Abcam | ab177487 | 1:300 | 1:300 |
| **NeuroD1** | MC-Mouse | Abcam | ab60704 | 1:1000 | 1:1000 |
| **S100B** | MC-Mouse | Sigma | SAB4200671 | 1:500 | - |
| **Smi31** | MC-Mouse | BioLegend | 801601 | 1:500 | - |
| **Sox2** | PC-Rabbit | Abcam | ab97959 | 1:500 | - |
| **Stem121** | MC-Mouse | Takara | Y40410 | - | 1:400 |
| **Synapsin I** | PC-Rabbit | Abcam | ab8 | 1:1000 | 1:1000 |
| **TH** | PC-Rabbit | Merk | AB152 | 1:500 | - |
| **Tuj1** | MC-Mouse | Covance | MMS-435P | 1:500 | 1:1000 |
| **vGlut1** | PC-Guinea Pig | Synaptic Systems | 135 304 | 1:2000 | - |
| **Vimentin** | PC-Chicken | Abcam | ab24525 | 1:1000 | - |

Table 2. Reprogramming of hmOEG to neurons with NEUROD1 + SM

| SM (µM) | | | | | | | | | | | Removal SM (DPI) | DPI | Result |
|---|---|---|---|---|---|---|---|---|---|---|---|---|---|
| FSK | CHIR | SAG | P | ISX9 | SB | DAPT | LDN | IBET | VPA | Y27 | | | |
| 20 | | | | | | | | | | | 14 / No | 21 | Cell death |
| 10 | 1,5 | | | | | | | | | | 14 / No | 21 | NI |
| 10 | 3 | | | | | | | | | | 14 / No | 21 | NI |
| 10 | | 0,1 | 0,1 | | | | | | | | 14 / No | 21 | NI |
| 10 | | | | 10 | | | | | | | 14 / No | 21 | NI |
| 10 | | | | | 5 | | | | | | 14 / No | 21 | NI |
| 10 | | | | | | 5 | | | | | 14 / No | 21 | NI |
| 10 | | | | | | | 0,25 | | | | 14 / No | 21 | NI |
| 10 | | | | | | | | 2 | | | 14 / No | 21 | NI |
| 10 | | | | | | | | | 500 | | 14 / No | 21 | NI |
| 10 | | | | | | | | | | 1 | No | 21 | NI |
| | | | | | 5 | | 0,25 | | | | 14 / No | 21 | NI |

**Table 3. Reprogramming hmOEG to neurons with SM**

| SM (µM) | | | | | | | | | | | Removal SM (DPI) | DPI | Result |
|---|---|---|---|---|---|---|---|---|---|---|---|---|---|
| **FSK** | **CHIR** | **SAG** | **P** | **ISX9** | **SB** | **DAPT** | **LDN** | **IBET** | **VPA** | **Y27** | | | |
| 10 | | | | | | | | | | | No | 21 | NR |
| 10 | 3 | | | | | | | | | | No | 21 | NR |
| 10 | | 0,1 | 0,1 | | | | | | | | No | 21 | NR |
| 10 | | | | 10 | | | | | | | No | 21 | NR |
| 10 | | | | | 5 | | | | | | No | 21 | NR |
| 10 | | | | | | 5 | | | | | No | 21 | NR |
| 10 | | | | | | | 0,25 | | | | No | 21 | NR |
| 10 | | | | | | | | 2 | | | No | 21 | NR |
| 10 | 3 | | | 10 | 5 | | | | | | No | 21 | NR, cell death |
| 10 | 3 | | | 10 | | 5 | | | | | No | 21 | NR, cell death |
| 10 | | | | | 5 | 5 | 0,25 | | | | No | 21 | NR |
| | | | | | 5 | | 0,25 | | | | No | 21 | NR |

**Table 4. Reprogramming hmOEG to neurons with SM combinations from the literature.**

| Reference (* modified protocol) | NEUROD1 | SM (d = dpi with SM) | | | | | | | | | | | DPI | Result |
|---|---|---|---|---|---|---|---|---|---|---|---|---|---|---|
| | | **FSK** | **CHIR** | **SAG, P** | **ISX9** | **SB** | **DAPT** | **LDN** | **IBET** | **VPA** | **Y27** | **AA** | | |
| (Gao *et al.*, 2017) | w / w.o. | d1-30 | d1-20 | | d1-20 | d1-20 | | | d1-30 | d1-20 | | d1-30 | 30 | NI/NR |
| (Gao *et al.*, 2017) * | w / w.o. | d1-30 | d1-8 | | d1-8 | d1-8 | | | d1-30 | d1-8 | | d1-30 | 30 | NI/NR |
| (Zhang *et al.*, 2015) | w / w.o. | | d3-7 | d7-9 | | d1-3 | d3-7 | d1-3 | | d3-5 | d1-9 | | 30 | NI/NR |
| (Yin *et al.*, 2019) | w / w.o. | | d2-6 | | | d1-2 | d2-6 | d1-2 | | | d6-30 | d6-30 | 30 | NI/NR |
| (Yin *et al.*, 2019) * | w / w.o. | | d1-6 | | | d1-6 | d1-6 | d1-6 | | | d6-30 | d6-30 | 30 | NI/NR |

SM: small molecules. DPI: days post induction. FSK: forskolin (PKA activator in cAMP signaling pathway). CHIR: CHIR99021(GSK3 inhibitor). SAG: Smoothened agonist (Sonic Hedgehog-SHH- activator). P: purmorphamine (SHH activator). ISX9: Isoxazole9 (inducer of adult neural stem cell differentiation). SB: SB431542 (TGFb receptor inhibitor). DAPT (gamma-secretase inhibitor). LDN: LDN193189 (BMP pathway inhibitor). IBET: IBET-151(BET bromodomain inhibitor). VPA: valproic acid (HDAC inhibitor). Y27: Y27632 (ROCK inhibitor). AA: ascorbic acid (antioxidant). NI: no improvement in reprogramming. NR: no reprogramming.
