## Supplemental Figure 1 for "GENERATION OF FUNCTIONAL NEURONS FROM ADULT HUMAN MUCOSAL OLFACTORY ENSHEATHING GLIA BY DIRECT LINEAGE CONVERSION"

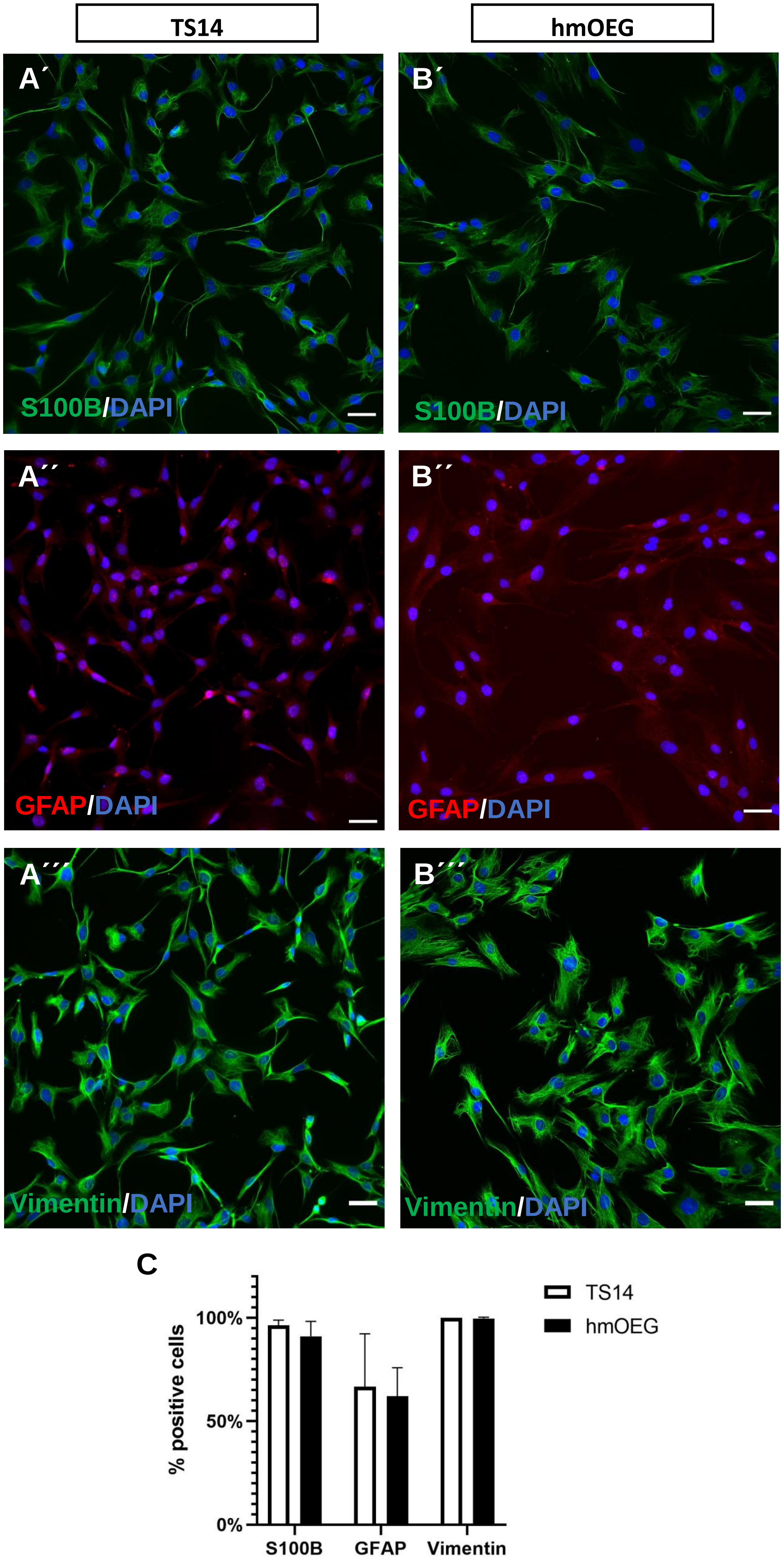

TS14
hmOEG
A´
S100B/DAPI
B´
S100B/DAPI
A´´
GFAP/DAPI
B´´
GFAP/DAPI
A´´´
Vimentin/DAPI
B´´´
Vimentin/DAPI
C
% positive cells
100%
50%
0%
S100B
GFAP
Vimentin
TS14
hmOEG

**Figure S1. Characterization of hmOEG cell line: analysis of OEG markers.** Expression analysis of glial markers was performed by immunofluorescence in a primary cell line of human mucosa olfactory ensheathing glia (hmOEG); an immortalized OEG cell line (TS14) was used as a positive control. Glial markers S100B (green) and vimentin (green) were expressed in TS14 (A′, A′′′) and hmOEG (B′, B′′′), while GFAP (red) was also expressed in both OEG cell types (A′′, B′′) but with a more diffused pattern. Nuclei were stained with DAPI (blue). The histograms (C) represent the mean±SD of the quantifications of triplicates. The statistical test applied was the Student's t-test (n=3, 25 fields were analyzed per experiment). Scale bar: 50 µm.
