## Supplemental Figure 2 for "GENERATION OF FUNCTIONAL NEURONS FROM ADULT HUMAN MUCOSAL OLFACTORY ENSHEATHING GLIA BY DIRECT LINEAGE CONVERSION"

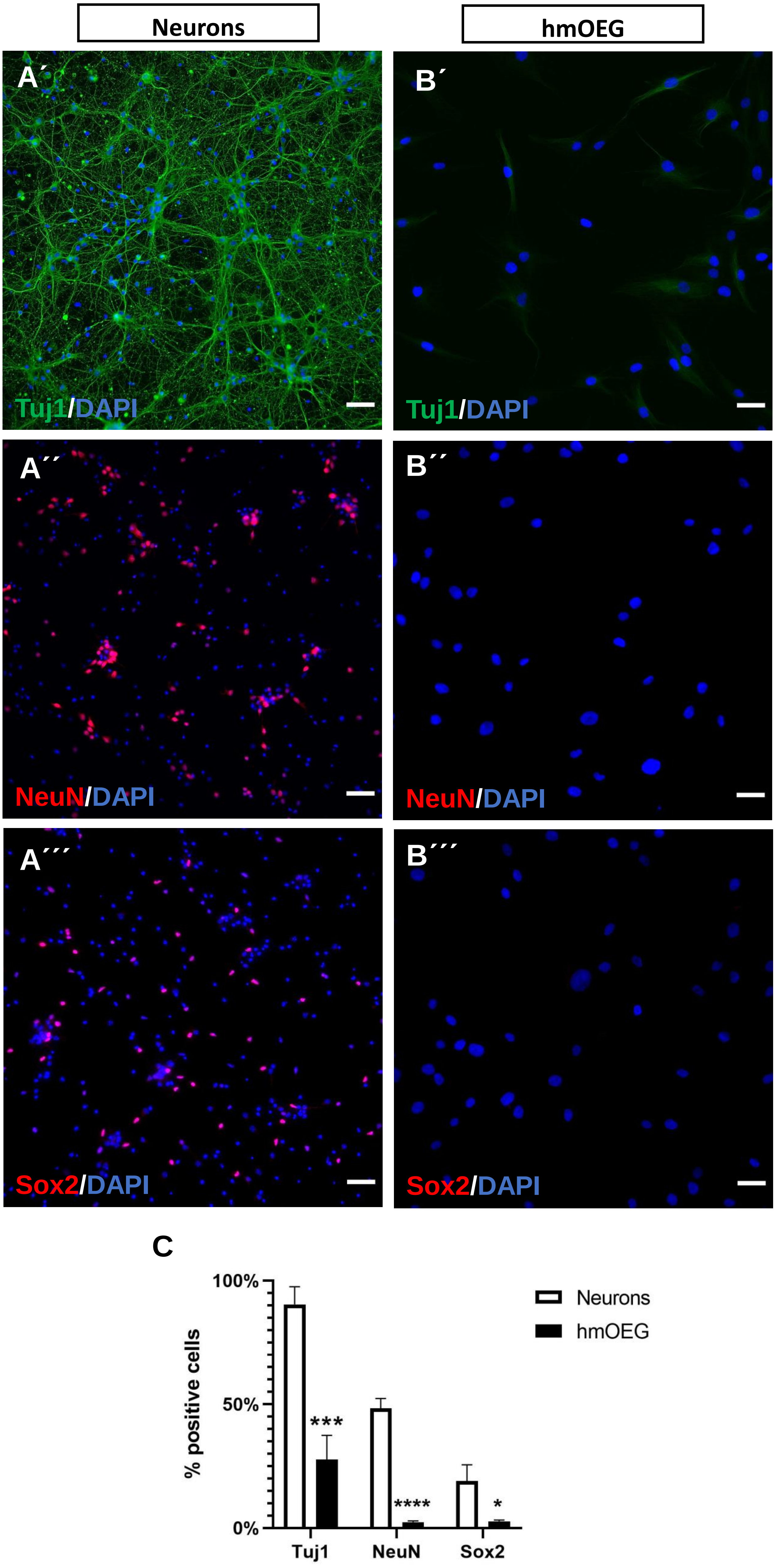
Neurons
hmOEG
A´
B´
Tuj1/DAPI
Tuj1/DAPI
A´´
B´´
NeuN/DAPI
NeuN/DAPI
A´´´
B´´´
Sox2/DAPI
Sox2/DAPI
C
100%
50%
0%
% positive cells
***
****
*
Tuj1
NeuN
Sox2
Neurons
hmOEG

**Figure S2. Characterization of hmOEG cell line: analysis of neuronal markers.** Expression analysis of neuronal markers was performed by immunofluorescence in a primary cell line of human mucosa olfactory ensheathing glia (hmOEG); a neuronal primary culture from rat embryonic cortex (E18) was used as a positive control. Proneural gene NeuN expression (red) was absent from hmOEG, while neuronal culture stained positively for this marker (A′′, B′′). However, Tuj1 (green) not only stained embryonic neurons but was also detected in hmOEG (A′, B′). hmOEG population was not contaminated with neural precursors as SOX2 positive staining (red) was not detected in the hmOEG culture, while neural progenitor cells from rat embryonic cortex expressed this neural precursor marker (A′′′, B′′′). Nuclei were stained with DAPI (blue). The histogram (C) represent the mean±SD of the quantifications of triplicates. The statistical test applied was the Student's t-test (n=3, 25 fields were analysed per experiment). Scale bar: 50 µm.
