## Supplemental Figure 3 for "GENERATION OF FUNCTIONAL NEURONS FROM ADULT HUMAN MUCOSAL OLFACTORY ENSHEATHING GLIA BY DIRECT LINEAGE CONVERSION"

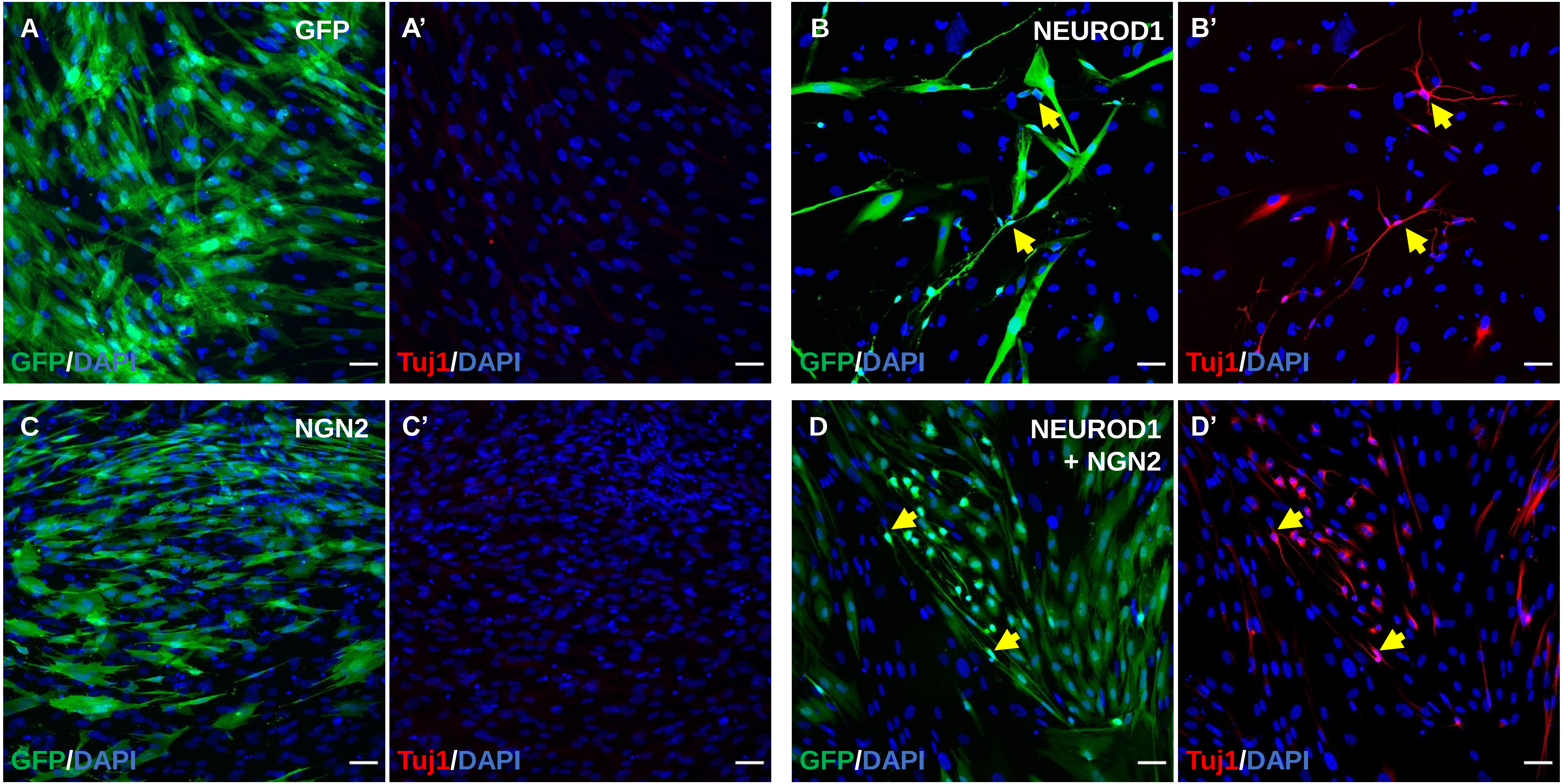
A
GFP
GFP/DAPI
A’
Tuj1/DAPI
B
NEUROD1
GFP/DAPI
B’
Tuj1/DAPI
C
NGN2
GFP/DAPI
C’
Tuj1/DAPI
D
NEUROD1
+ NGN2
GFP/DAPI
D’
Tuj1/DAPI

**Figure S3. Screening of transcription factors to convert hmOEG to induced neurons (hmOEG-iNs).** Representative images of the expression of the neuronal marker Tuj1, 30 dpi in hmOEG infected with GFP alone (A, A´), NEUROD1 (B, B´), NEUROG2 (C, C´) or in combination (D, D´). Yellow arrows indicate GFP+ cells expressing Tuj1. Scale bar: 50 µm. dpi: days post-infection. hmOEG: human mucosa adult olfactory ensheathing glia.
