## Supplemental Figure 4 for "GENERATION OF FUNCTIONAL NEURONS FROM ADULT HUMAN MUCOSAL OLFACTORY ENSHEATHING GLIA BY DIRECT LINEAGE CONVERSION"

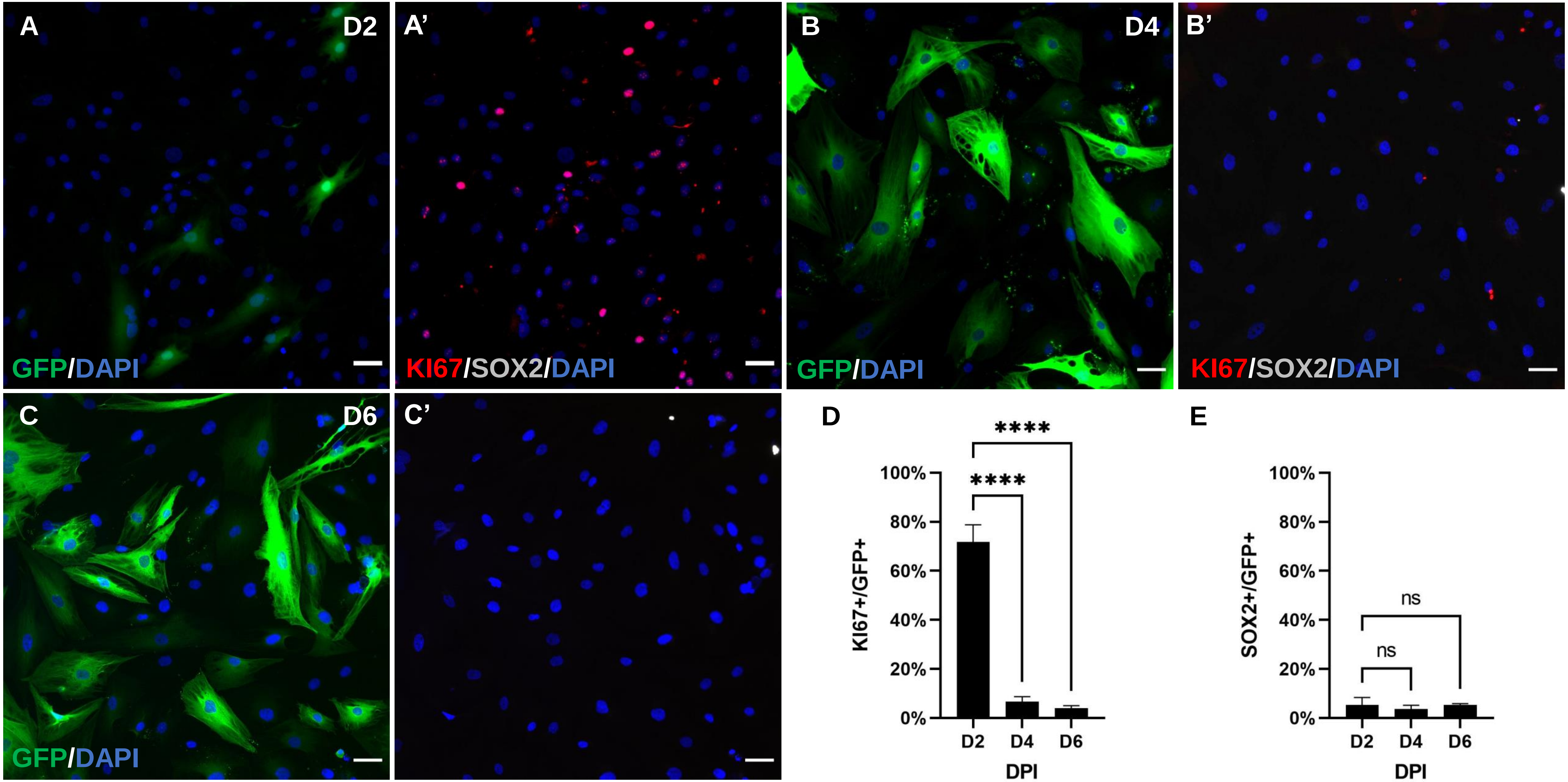
A
D2
GFP/DAPI
A'
KI67/SOX2/DAPI
B
D4
GFP/DAPI
B'
KI67/SOX2/DAPI
C
D6
GFP/DAPI
C'
D
****
****
100%
80%
60%
40%
20%
0%
KI67+/GFP+
D2
D4
D6
DPI
E
ns
ns
100%
80%
60%
40%
20%
0%
SOX2+/GFP+
D2
D4
D6
DPI

**Figure S4: NEUROD1-hmOEG does not transition to a neural stem cell-like state before differentiating into iNs.** Representative images of SOX2 (white) and KI67 (red) expression at 2, 4 and 6 dpi. NEUROD1 infected hmOEG (GFP+ cells) were in a proliferative stage at 2 dpi (expressing KI67), but at 4 and 6 dpi there was a significant drop in the percentage of cells expressing this marker. There was no significant variation over the days in the percentage of positive cells for SOX2. The histograms represent the mean±SD of the quantifications of triplicates of KI67 and SOX2. Statistical tests applied were One-way ANOVA and post-hoc Tukey test (****$p \leq 0.0001$ ; NS, not significant) for multiple comparisons between means (n=3, per experiment 10 fields were analysed). Scale bar: 50 µm. DPI: days post-infection.
