## Supplemental Figure 5 for "GENERATION OF FUNCTIONAL NEURONS FROM ADULT HUMAN MUCOSAL OLFACTORY ENSHEATHING GLIA BY DIRECT LINEAGE CONVERSION"

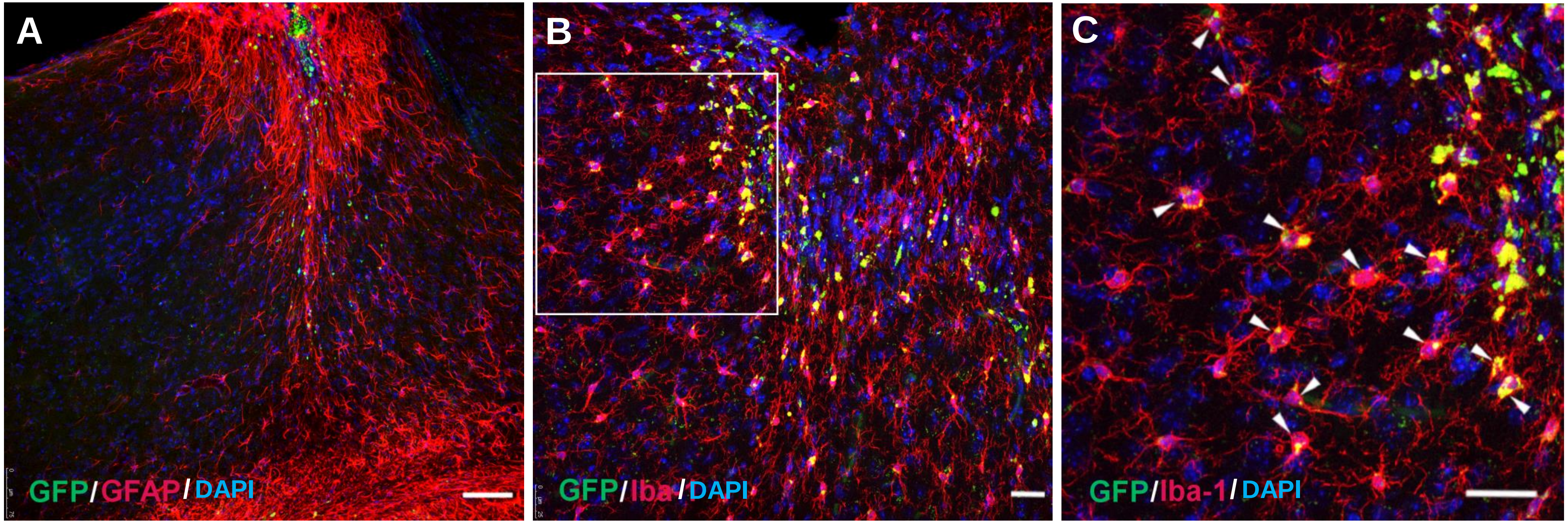
A
GFP/GFAP/DAPI
B
GFP/Iba/DAPI
C
GFP/Iba-1/DAPI

**Figure S5: Reactivity after hmOEG-iNs transplantation into the brain of NOD-SCID mice.** (A) Intense astroglial reactivity at the injection site revealed by immunostaining with the astrocytic marker GFAP (red). (B) Microglia localization at the injury site showed by immunostaining with Iba-1 microglial marker (red). GFP staining (green) identifies hmOEG cells. (C) Colocalization of GFP and Iba-1 staining (arrowheads) suggesting phagocytosis of hmOEG by microglia. Nuclei were stained with DAPI (blue). Scale bar: 75 µm (A) and 25 µm (B,C).
